# Investigating the Volume and Diversity of Data Needed for Generalizable Antibody-Antigen ∆∆G Prediction

**DOI:** 10.1101/2023.05.17.541222

**Authors:** Alissa M. Hummer, Constantin Schneider, Lewis Chinery, Charlotte M. Deane

## Abstract

Antibody-antigen binding affinity lies at the heart of therapeutic antibody development: efficacy is guided by specific binding and control of affinity. Here we present Graphinity, an equivariant graph neural network architecture built directly from antibody-antigen structures that achieves state-of-the-art performance on experimental ∆∆G prediction. However, our model, like previous methods, appears to be overtraining on the few hundred experimental data points available. To test if we could overcome this problem, we built a synthetic dataset of nearly 1 million FoldX-generated ∆∆G values. Graphinity achieved Pearson’s correlations nearing 0.9 and was robust to train-test cutoffs and noise on this dataset. The synthetic dataset also allowed us to investigate the role of dataset size and diversity in model performance. Our results indicate there is currently insufficient experimental data to accurately and robustly predict ∆∆G, with orders of magnitude more likely needed. Dataset size is not the only consideration – our tests demonstrate the importance of diversity. We also confirm that Graphinity can be used for experimental binding prediction by applying it to a dataset of *>*36,000 Trastuzumab variants.

---

Antibodies mediate their functions, both physiologically and therapeutically, by binding specifically to a target antigen. Controlling affinity is therefore the driving consideration in therapeutic antibody development when identifying, as well as optimizing, a lead candidate.

Many other properties, aside from affinity, often referred to collectively as developability, also play important roles. There have been substantial advances in recent years in using machine learning (ML) to predict such properties – from self-association [1] and humanness [2, 3] to polyreactivity and specificity [4, 5]. However, changes to the antibody sequence to improve these properties must not come at the cost of binding. Thus, therapeutic antibody development relies on solving a complex, multi-parameter optimization problem [6].

Experimental techniques for affinity quantification are typically slow and laborious [7]. A fast and accurate computational predictor of change in affinity would fill a need in the antibody design pipeline. Furthermore, computational approaches can, in principle, incorporate information from different predictors to simultaneously optimize multiple properties, while still controlling binding affinity.

*In silico* prediction of antibody-antigen affinity remains a challenge. Traditional affinity prediction tools, such as FoldX [8] and Rosetta [9], are based on physical equations and empirical measurements. These have proven effective for certain applications [10] but can be limited in speed and accuracy [9, 11]. In recent years, there has been a shift towards ML approaches, which can be divided into two main categories: sequence- and structure-based. Sequence-based methods have been successfully applied to predict affinity for a specific antigen in cases where a large amount of data is available [12, 13]. These methods are not broadly generalizable: the information they are trained on is antigen-specific and the models cannot be readily applied to another antigen without further training. Structure-based methods promise greater generalizability by aiming to capture the interaction patterns across many different antibody-antigen complexes. Current methods are trained on features derived from antibody-antigen complex structures, such as binding surface area, interatomic interactions and energy-based terms [11, 14, 15]. However, these methods appear to not predict well outside their training data [14, 16]. Additionally, they require the extraction of features, which can be slow and is subject to human bias.

Here we present Graphinity, an equivariant graph neural network (EGNN) architecture for predicting change in antibody-antigen binding affinity. Our deep learning models are built directly from protein complex structures, potentially enabling scalability and generalizability.

Graphinity achieved state-of-the-art performance for ∆∆G_prediction on single-point mutations from the experimental AB-Bind dataset [17], achieving test Pearson’s correlations of up to 0.80. However, further investigation indicated that this high performance stemmed from model overtraining and was not robust to train-test cutoffs, an observation which has been hinted at by results from previous methods [14, 16, 18, 19].

To examine if our architecture could be used to robustly predict change in binding affinity, we generated a large synthetic dataset of nearly 1 million ∆∆G values using FoldX [8]. We achieved Pearson’s correlations close to 0.9 on this dataset, which were robust to train-test sequence identity cutoffs and noise.

This far larger dataset allowed us to test the volume and type of data needed for a potentially generalizable antibody-antigen ∆∆G predictor. Investigating model performance with varying amounts of synthetic data demonstrated that there is currently insufficient experimental data to accurately predict ∆∆G, with orders of magnitude more likely to be needed. Our results also highlighted the importance of dataset diversity for model predictiveness.

We validated that Graphinity can learn not only the FoldX forcefield but also experimental binding affinity by adapting and successfully applying it to a dataset of *>*36,000 HER2-binding and non-binding Trastuzumab variants [12].

We have made the synthetic datasets and model code publicly available at https://github.com/oxpig/Graphinity.

## Results

### Graphinity model architecture

Graphinity takes structures of a wild-type (WT) and a mutant antibody-antigen complex as input, feeds the corresponding graph representations through a Siamese EGNN and predicts ∆∆G (Figure 1a). In the atomistic graphs, non-hydrogen atoms are represented as nodes and interactions between nodes less than 4 Å apart as edges. Graphs are limited to the neighborhood around the mutated site. The architecture is modular and easily adapted for regression/classification and for single-/multi-graph inputs. See Methods for full details.

**Figure 1:**
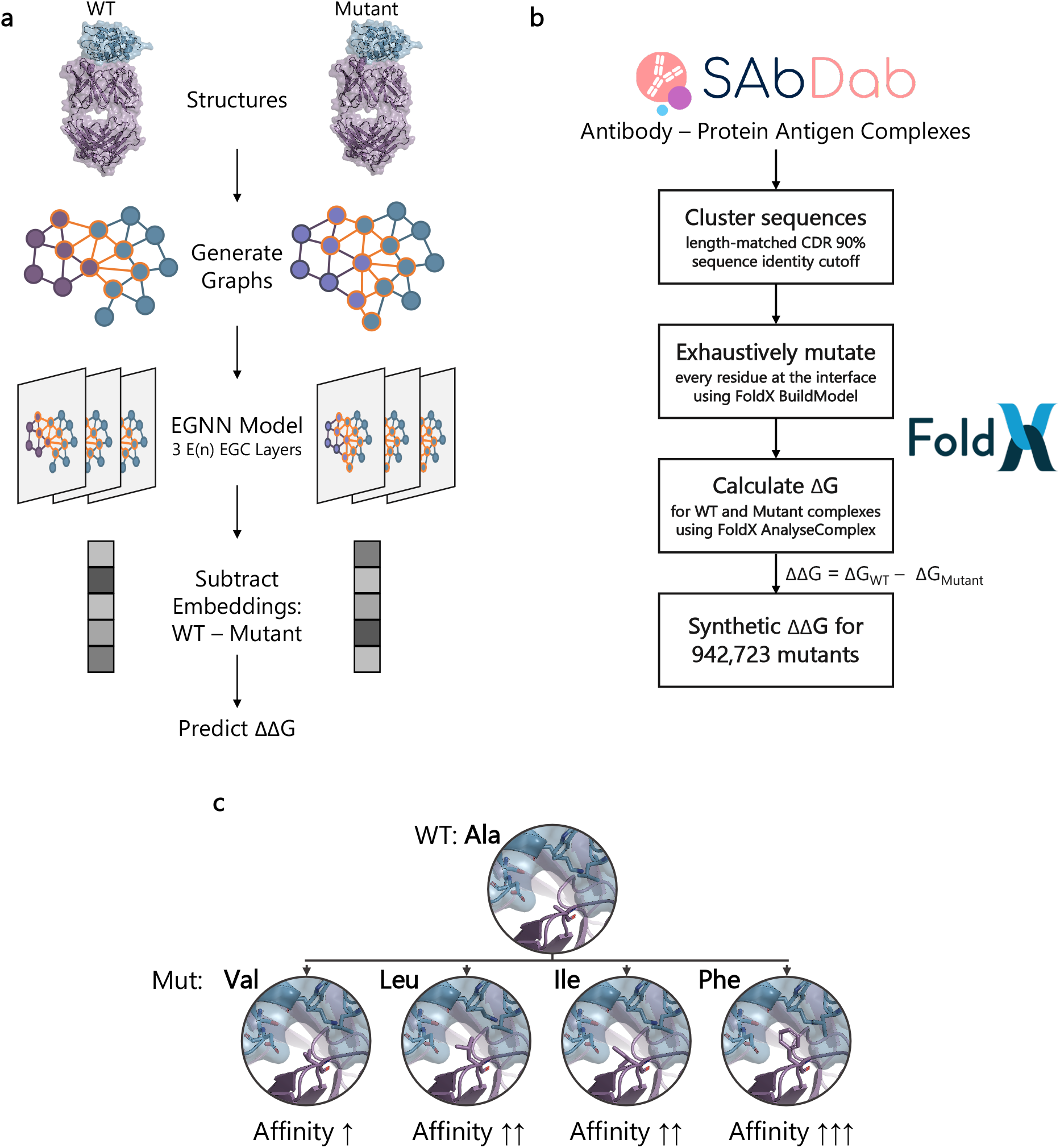
Graphinity architecture and synthetic dataset preparation. (a) The EGNN deep learning models are trained on graphs of 3D protein structure coordinates. The graphs are built from atoms in the neighborhood of the mutated site. Our model architecture consists of three E(n) Equivariant Graph Convolutional (EGC) layers [20], followed by a linear layer. For ∆∆G prediction, the embeddings generated from the E(n) EGC layers are subtracted from one another prior to passing through the linear layer. (b) The synthetic ∆∆G dataset was generated from structurally resolved complexes from SAbDab [21, 22]. We exhaustively mutated interface residues and applied FoldX AnalyseComplex to estimate the ∆G of the complexes [8]. (c) Example of ∆∆G data for a complex. PDB: 1XGP [23]; affinity values from SKEMPI 2.0 [24].

### Graphinity performance for predicting experimental ∆∆G

We applied Graphinity to the experimental ∆∆G dataset from AB-Bind [17], which contains 645 single-point mutations from 29 complexes and will from here on be referred to as Experimental ∆∆G 645 (see Supplementary Table 1 for a description of this dataset, Supplementary Figure 1a for the ∆∆G distribution and Figure 1c for an example of ∆∆G data). We considered hypothetical reverse mutations in the training and validation data (Experimental_∆∆G_645 + Reverse Mutations), in which the WT and mutant structures were swapped and the corresponding label was given the opposite sign, an approach taken by others [11, 15]. Additionally, the AB-Bind dataset contains non-binder mutations with ∆∆G values arbitrarily set to −8 kcal/mol. We applied our model to datasets including and excluding these non-binders (Experimental_∆∆G_645 ±Non-Binders).

Our model achieved Pearson’s correlations of up to 0.80 on 10-fold cross-validation (Figure 2a), similar in performance to existing methods which report correlations of up to 0.76 [14, 15]. However, delving into the robustness of the model – by imposing sequence identity cutoffs between folds – indicated that these high correlations were the result of overtraining as opposed to true learning (Figure 2b). When we imposed a 100% length-matched CDR sequence identity cutoff, ensuring that mutations from the same complex cannot be in both the training and test dataset, the Pearson’s correlations decreased by an average of 63%. The results were also highly sensitive to the inclusion of non-binders (Figure 2b) and, across all train-test cutoffs, there was substantial variation in the Pearson’s correlation across different folds (Supplementary Figure 2). Poor model robustness on experimental ∆∆G prediction has been found for previous approaches [14, 16, 18, 19].

**Figure 2:**
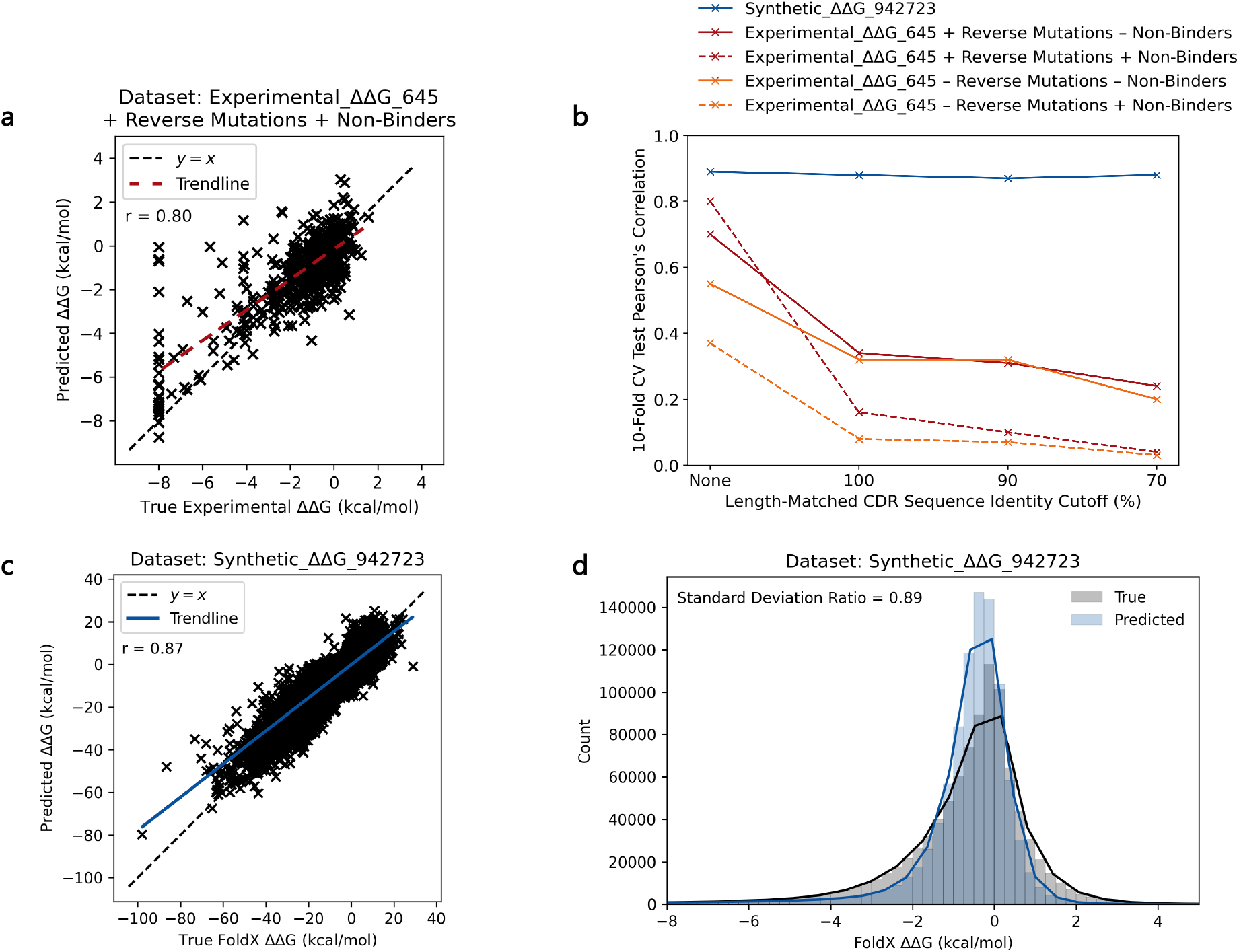
Graphinity model performance for ∆∆G Prediction. (a) Graphinity achieves a Pearson’s correlation of 0.80 on the Experimental_∆∆G_645 + Reverse Mutations + Non-Binders dataset, with a random train-validation-test split. Reverse mutations were used for training/validation only and were not included in the test dataset. An ensemble of 10 models was trained for 500 epochs with 10-fold cross-validation (CV) on the datasets. The trendline, shown in red, is a least squares polynomial fit. (b) Model performance decreases substantially for experimental ∆∆G prediction when a sequence identity cutoff is applied to the train-validation-test split. The models trained and tested on the large synthetic dataset generated using FoldX [8] (blue) are far more robust. This figure is included with error bars representing the standard deviation across the 10 folds in Supplementary Figure 2. (c) Graphinity achieves a Pearson’s correlation of 0.87 on the Synthetic_∆∆G_942723 dataset with a 90% length-matched CDR sequence identity cutoff applied for the train-validation-test split. An ensemble of 10 models was trained for 10 epochs with 10-fold cross-validation. The trendline, shown in blue, is a least squares polynomial fit. (d) Histograms of the true and predicted FoldX ∆∆G values (x-axis limited to −8 to +5 kcal/mol for clarity), shown in (c), demonstrate good agreement between the two sets of values. The solid lines are kernel density estimates (KDEs).

Tests of existing methods for antibody-antigen ∆∆G prediction have not imposed any cut-off between cross-validation folds, with the exception of leave-one-complex-out cross-validation, in which mutations from the same PDB cannot be in both the training and test set (although mutations from identical or closely related complexes in separate PDBs could be) [14, 15]. For one method, TopNetTree, this leave-one-complex-out test caused a drop in the average Pearson’s correlation to 0.17 [14]. For another method, mCSM-AB2, only a minor drop in performance was reported but they appear to include hypothetical reverse mutations in their test data [15].

Although widely used, the AB-Bind dataset suffers from several limitations, including that five of the included complexes do not contain an antibody, despite the dataset being described as an “antibody binding mutational database” [17] (more details in Methods). We therefore propose a new experimental antibody-antigen single-point mutation ∆∆G dataset (Experimental_∆∆G_608, Supplementary Table 1, Supplementary Figure 1b), consisting of 608 mutations filtered from the SKEMPI 2.0 database [8], for model benchmarking. Although this dataset has fewer single-point mutations, these mutations come from a slightly larger number of complexes (33). The performance of Graphinity on this dataset is similar to that for the Experimental_∆∆G_645 dataset (Supplementary Figure 3a): model correlation is high when the data is split randomly and with reverse mutations, but once again is not robust to train-test CDR sequence identity cutoffs.

On this more rigorous dataset, we also investigated the role of model architecture. We built a tree-based model from features (more details in Methods) derived from the WT and mutant complex structures, similar to the method employed by the mCSM-based models [11, 15]. This different model architecture gave similar correlations but also suffered from overtraining (Supplementary Figure 3b), suggesting that the problem lies in the data.

### Using a synthetic dataset of ∼1 million mutations

The poor robustness of model performance on the limited experimental data led us to investigate how well ∆∆G could be predicted if more data was available. We generated a synthetic dataset of nearly 1 million ∆∆G data points (Synthetic_∆∆G_942723, Supplementary Table 1, Supplementary Figure 1c) by exhaustively mutating the interfaces of structurally-resolved complexes from the Structural Antibody Database (SAbDab) [21, 22] using FoldX [8] (Figure 1b). This synthetic dataset will not completely mimic the complexity of true ∆∆G values. The Pearson’s correlation between FoldX predictions and experimental values is 0.34 for the AB-Bind dataset [17]. The accuracy is higher for mutations with a larger effect on binding affinity though. The area under the receiver operating characteristic curve (ROC AUC) in predicting whether a mutation is stabilizing or not is 0.87 for mutations with an absolute value greater than 1 kcal/mol [17], suggesting that this data does contain some of the characteristics of experimental values.

On this synthetic dataset, Graphinity achieved a test Pearson’s correlation of 0.87 with 10-fold cross-validation and a 90% length-matched CDR sequence identity cutoff imposed between folds (Figure 2c). Training the model for longer (100 epochs, as opposed to 10) improved the correlation slightly, to 0.91 on a single fold, but we did not explore this further as it was computationally costly to run. Graphinity substantially outperformed a simple baseline for predicting ∆∆G: the correlation between the change in number of contacts between the WT and mutant structure (4 Å interaction distance cutoff) and the synthetic ∆∆G is 0.42, less than half the correlation our EGNN model achieves.

The performance of Graphinity was robust to train-validation-test sequence identity cutoffs (Figure 2b; Figure 4a). The most stringent split, a length-matched CDR sequence identity cutoff of 70% plus an antigen sequence identity cutoff of 70%, maintained a Pearson’s correlation above 0.87. For reference, 88% of antibody therapeutics share at least 70% heavy and light chain CDR sequence identity with a natural antibody sequence [25].

Another way to assess model performance is with the Spearman’s rank correlation. This value (ca. 0.6) was lower than the Pearson’s correlation for our models. This appears to be due, in large part, to the very high density of ∆∆G values close to 0, which the EGNN did not always rank in the correct order. The Spearman’s rank correlation rose to ca. 0.7 when values between −1 and +1 kcal/mol were excluded. FoldX is known to be less accurate at predicting the ∆∆G values for mutations with only a small effect on binding affinity [17] and therefore there may be less signal in the data in this region.

A recent study found that FoldX accuracy was higher for mutations that are observed naturally, with an 11% decrease in incorrectly predicting a mutation to be stabilizing [26]. We explored the performance of our model on a test dataset limited to such ‘evolutionarily grounded’ mutations, as defined in [26], and found that the Pearson’s correlation was stable at 0.89 (for more details see Supplementary Information).

We also investigated model performance with different graph inputs – of the full interface rather than just the mutation site neighborhood, reflecting the input for potential multi-point mutation data – and found that performance was maintained (Pearson’s correlation = 0.85 on held-out test data, 90% length-matched CDR sequence identity cutoff).

These results serve as a proof of concept that ∆∆G can be accurately predicted when sufficient data is available.

### Considerations for generating experimental ∆∆G datasets

Having demonstrated the potential of our EGNN architecture for predicting ∆∆G when input data is abundant, we next attempted to quantify the amount of data that will be required for the accurate prediction of experimental values. We built models with varying training plus validation dataset sizes (datasets Synthetic_∆∆G_{580-450000}, Supplementary Table 1) and applied them to a test set of 94,126 mutations (90% length-matched CDR sequence identity cutoff). We found that test Pearson’s correlations only began to plateau, reaching 0.85, for models trained with at least 90,000 mutations (Figure 3a).

**Figure 3:**
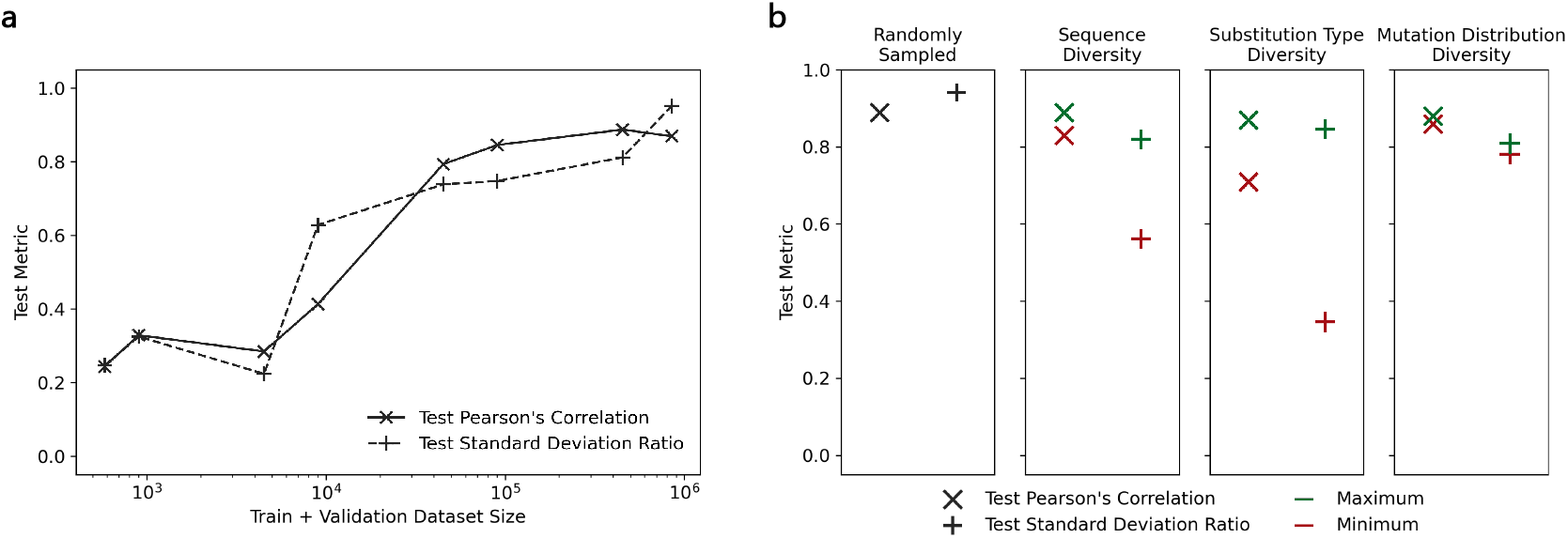
Considerations for experimental ∆∆G dataset generation, with respect to ML predictiveness. (a) Graphinity performance is highly dependent on the size of the dataset, only achieving correlations of ca. 0.85 for datasets with 90,000 mutations and more. While the test Pearson’s correlation does not increase substantially past this point, the distribution of predicted ∆∆G values continues to widen, becoming more similar to the distribution of the true synthetic ∆∆G values, representing greater model predictiveness (standard deviation ratio). Datasets used: Synthetic_∆∆G_{580-450000} (Supplementary Table 1). (b) Investigating the influence of dataset diversity on model performance, we considered diversity in antibody CDR sequence identity, amino acid substitution type frequency and the distribution of mutated positions in the complex. Model performance is substantially lower for the sequence and substitution type minimum-diversity datasets, particularly as seen through the standard deviation ratio. Datasets used: Synthetic_∆∆G_100000 randomly sampled, Synthetic_∆∆G_100000_{sequence/substitution type/substitution_distribution}_{min/max}(Supplementary Table 1).

Upon comparing the distributions of the predicted and true values, we observed that the models built from smaller datasets often regressed towards the mean and achieved high correlations despite predictions not covering the full range of true values. To quantify this effect, we calculated the standard deviation ratio, the relative ratios of the standard deviations of the true and predicted values. The standard deviation ratio does not plateau at any stage and only exceeds 0.8 with a dataset size of 450,000 mutations (Figure 3a).

Diversity is a known important characteristic of any dataset used for model training. We evaluated the role of dataset diversity using three metrics: the diversity of antibody sequences, amino acid substitution types and structural distribution of mutations in the interface (for more details see Methods). We constructed training and validation datasets to minimize and maximize each respective metric (Synthetic_∆∆G_100000_{sequence/substitution type/substitution distribution}_{min/max}, Supplementary Table 1). For example, the Synthetic_∆∆G_100000_sequence_min_training dataset contained mutations from 75 antibody-antigen complexes, while the corresponding maximum-diversity dataset contained mutations from 1177 complexes. All models built from these datasets were evaluated on the same test data, consisting of 10,000 mutations (Supplementary Table 1). A 90% length-matched CDR sequence identity cutoff was imposed between all the training, validation and test datasets.

The distribution of mutations in the interface had only a marginal effect, which may be explained by the input graphs, which represent only the neighborhood of the mutated site. However, we found that sequence and substitution type diversity impacted model performance, particularly the test standard deviation ratio (Figure 3b). The minimum sequence and substitution type diversity datasets achieved 31% and 59% lower standard deviation ratios than the corresponding maximum diversity datasets, respectively.

### Graphinity is robust to noise on large synthetic ∆∆G dataset

Experimental ∆∆G data is noisy, particularly if acquired from different experimental setups and/or labs [24]. We therefore explored the robustness of Graphinity to noise by perturbing the training and validation datasets of Synthetic_∆∆G_942723 in two ways: (1) shuffling the affinity labels corresponding with mutations (Synthetic_∆∆G_942723_shuffled) and (2) adding Gaussian-distributed random noise to the labels (Synthetic_∆∆G_942723_gaussian_noise).

The Pearson’s correlations on held-out test sets remained remarkably constant, at approximately 0.85 for datasets with 0-60% shuffled labels (Figure 4b). However, upon analyzing the relative distributions of the predicted and true FoldX ∆∆G values, we found that the model lost predictiveness with increased shuffling: the predicted values began to fall in increasingly narrow distributions as compared to the true spread of ∆∆G (Figure 4b). These results underscore the importance of looking beyond the traditional evaluation metric of Pearson’s correlation and also assessing the standard deviation ratio. Model performance was 0 when 100% of the labels were shuffled, supporting that, while the FoldX-generated values are not as accurate as experimental data, there is true signal that can be learned from the input complex structures.

**Figure 4:**
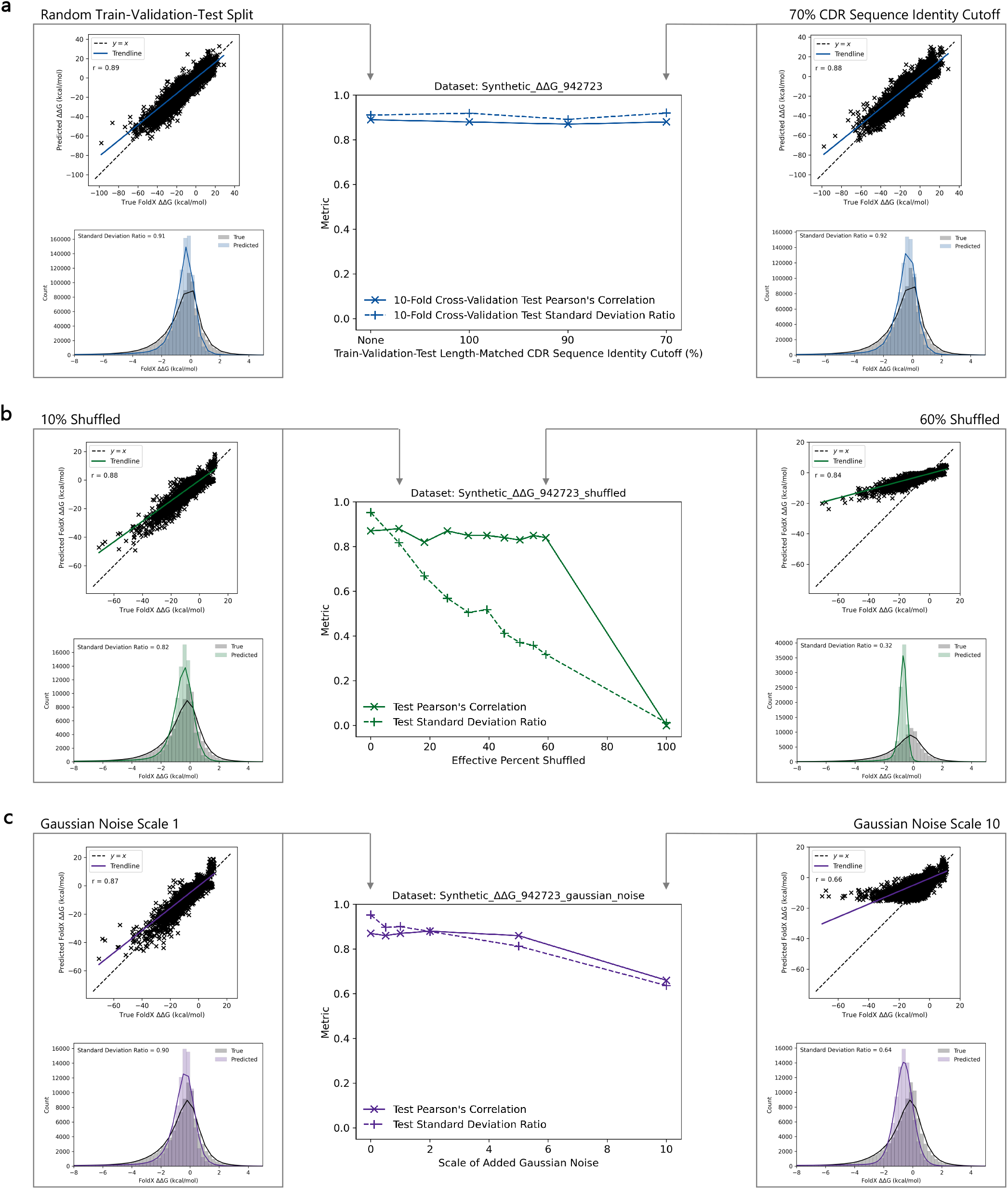
Graphinity is robust to train-validation-test cutoffs and noise on synthetic data. (a) Train-validation-test cutoffs: Graphinity performance is maintained across length-matched CDR sequence identity cutoffs applied when splitting the train, validation and test datasets. The synthetic dataset was already filtered such that no complex had more than 90% CDR sequence identity with any other complex and as such, the 100% and 90% cutoffs are functionally identical (although these were sampled from the full dataset separately). (b) Shuffling (Synthetic_∆∆G_942723_shuffled): Model test Pearson’s correlations stay relatively consistent as the percentage of labels which are shuffled (i.e. incorrect) increases. However, the distribution of predicted values narrows considerably at higher shuffling percentages, indicating that the model is losing predictiveness. (c) Gaussian noise (Synthetic_∆∆G_942723_gaussian_noise): Graphinity is robust to random noise added to the training and validation dataset, maintaining test Pearson’s correlations and standard deviation ratios above 0.8 up to a Gaussian noise scale of 5. Results are shown for 10-fold cross-validation in (a) and for a single fold, held-out test set in (b-c). For all histograms, the x-axes were limited to −8 to +5 kcal/mol for clarity and the solid lines are KDEs.

There are 82 duplicated antibody-antigen single-point mutations in SKEMPI 2.0 [8] which did not result in non-binders or have imprecisely measured affinity. Across these, the average ∆∆G standard deviation between duplicates is 0.19 kcal/mol and the maximum 0.90 kcal/mol. Graphinity maintained Pearson’s correlations and standard deviation ratios above 0.8 with added noise in this range, and indeed up to a Gaussian noise scale of 5 (Figure 4c).

### Performance by amino acid substitution

We further investigated how our model performs for specific amino acid substitutions (e.g. Arg to Lys). The ∆∆G values varied widely for a specific substitution, with standard deviations ranging from 0.5 to 10.6 kcal/mol (Supplementary Figure 4c). The pattern of the mean ∆∆G values and corresponding standard deviations closely matched between experimental and predicted values (Supplementary Figure 4), suggesting the model is learning the structural context of the mutations rather than just the average value for the mutation. If the predicted values were set as the average ∆∆G value for the specific substitution, the Pearson’s correlation would be just 0.35 as compared with the trained model’s performance of 0.87.

We also trained models on datasets limited to each substitution type to explore whether Graphinity could learn the effect of a substitution better when trained only on data for this substitution. However, model performance for a specific substitution decreased as compared to the model trained on the full dataset (Supplementary Figure 5). Performance could be rescued, reaching or exceeding that of the model trained on the full dataset, by initializing with weights from the full model (Supplementary Figure 5).

### Validation on experimental binding dataset

To test whether Graphinity can learn the distribution of experimental data, not just FoldX predictions, we adapted and applied our architecture to a dataset of 36,391 CDRH3 variants of Trastuzumab [12]. The variants are classified as binders or non-binders for the antigen, HER2. While a single-antigen task is not necessarily the intended aim of the Graphinity architecture, this dataset was sufficiently large that we would expect prediction to be successful.

We modeled the structures of the CDRH3 variants, mutated at up to 10 positions, using FoldX BuildModel [8]. Although this approach is unlikely to capture the true structural effect of the mutations, as FoldX does not model changes to the backbone [27], it is fast and allows us to avoid docking by starting from a structure of a bound complex. We modified Graphinity to take only one graph as input (of the variant, rather than WT and mutant structures), to build the input graphs around the mutated CDRH3 residues and their surrounding neighborhood (rather than a single residue position) and for a classification task (for details see Methods).

Our model learned to separate the binding and non-binding variants, achieving a ROC AUC of 0.88 and average precision (AP) of 0.77 (Supplementary Figure 6). This performance is close to that of the sequence-based convolutional neural network (CNN) reported by the authors of the study (ROC AUC = 0.91, AP = 0.83) [12]. Furthermore, our performance was robust to train-validation-test cutoffs with ROC AUC values maintained above 0.83 when V- and J-gene clonotype plus CDRH3 sequence identity cutoffs (90%, 70%) were applied (Supplementary Figure 6b).

## Discussion

Antigen binding affinity, essential to the function and efficacy of an antibody, is complex and challenging to predict computationally. Graphinity is built directly from the coordinates of antibody-antigen structures and does not rely on featurization, which is slow and may miss information that could be important for predicting affinity. We applied this architecture to ∆∆G prediction on both the limited available experimental data and a large constructed synthetic dataset. Graphinity achieved state-of-the-art performance on the experimental data from the AB-Bind database [17]. However, the high correlations were the result of overtraining, as has been found across existing ML methods for ∆∆G prediction [14, 16, 18, 19]. Overtraining is also gaining increased recognition in related fields, such as protein-protein interaction prediction [28]. To prevent information leakage, effective sequence identity cutoffs between train and test datasets are essential.

To test whether affinity could be accurately and robustly predicted, we applied Graphinity to a synthetic dataset of nearly 1 million mutants [8]. Test Pearson’s correlations on this dataset neared and the model generalized well beyond its training data, with performance being maintained with stringent sequenc0.9 e identity cutoffs for both antibody and antigen between the train, validation and test datasets.

Performance was also robust to levels of noise that have been observed in experimental data. Applying Graphinity to noisy data emphasized the importance of going beyond the test Pearson’s correlation when evaluating a model. A high correlation can be achieved when the model regresses towards the mean, predicting values in only a small range and with a trendline much flatter than *y* = *x*. The standard deviation ratio, a metric comparing the relative distributions of the true and predicted values, exposes poor predictiveness by identifying when predicted values cover only a fraction of the true ∆∆G distribution.

Our results on the synthetic data must be considered in light of the source of the data. The synthetic data points were all produced by the same software and are thus expected to be more self-consistent and less noisy than experimental data. The synthetic values may also follow a different distribution to the true values. However, FoldX can accurately predict whether mutations with a substantial effect on binding affinity will be stabilizing or destabilizing, suggesting there is signal in this dataset [17].

To test if the Graphinity architecture can also learn the distribution of experimental data, not just the FoldX forcefield, we applied it to an experimental dataset of 36,391 Trastuzumab variants. Graphinity separated binding from non-binding variants with a ROC AUC similar to that achieved by a CNN trained on the variant sequences. The EGNN architecture offers further benefits over the CNN, most notably the potential for generalizability to different antibody-antigen complexes. This application also highlights the modularity of our architecture: Graphinity can be applied for regression and classification, single- and multi-point mutations, as well as affinity and change in affinity prediction.

The success of our model on large datasets lends support to the idea that the major challenge with experimental ∆∆G prediction lies in the availability of experimental data rather than the model architecture. We explored the amount of data that would be required for accurate and generalizable prediction of experimental ∆∆G using the synthetic dataset. Our results suggest that there is currently vastly insufficient data available and orders of magnitude more, tens to hundreds of thousands of data points, will likely be needed. There is potential for limitations in data to be compensated for, to some extent, by machine learning know-how such as by identifying model architectures that require less data, using stratified sampling or by transfer learning from a related data-rich task or from synthetic data. Future model design could also be augmented by considering physiological features which are typically ignored in current methods, such as water molecules and protein conformational flexibility.

In addition to dataset size, our results underscore the importance of dataset diversity, particularly with respect to antibody sequence identity and amino acid substitution type. Both of these diversity metrics are currently very limited in experimental data. For example, the antibody-antigen single-point mutations in SKEMPI 2.0 [24] derive from less than 50 complexes and are highly skewed in substitution type, with mutations to alanine making up over half of the dataset.

Our results highlight the need to move towards ‘machine learning-grade data’, where model development is considered in the data generation process.

## Methods

### Experimental ∆∆G data preparation

The AB-Bind dataset consists of 645 single-point mutations and ∆∆G measurements from 29 complexes. We downloaded this dataset, which was originally compiled by Sirin et al. [17], from TopNetTree [14]. We reversed the sign on the ∆∆G labels to reflect ∆∆G = ∆*G*_*W T*_ *−*∆*G*_*Mutant*_, as is done in [15] and our synthetic datasets. We ‘repaired’ the structures using FoldX (version 5) RepairPDB and modeled the mutations using FoldX BuildModel [8]. We refer to this dataset as Experimental ∆∆G 645 (Supplementary Table 1).

As in mCSM-AB2 [15], we generated reverse mutations by mutating the forward mutant model back to wild-type (WT) using FoldX BuildModel and setting the ∆∆G label to the negative value of the forward mutation (Experimental_∆∆G_645 + Reverse Mutations).

The Experimental_∆∆G_645 dataset has multiple limitations: 5 of the complexes do not contain an antibody (PDBs: 1AK4, 1FFW, 1JTG, 1KTZ, 3K2M), 27 of the mutations are non-binders whose change in binding affinity has arbitrarily been set to −8 kcal/mol and there are 3 duplicated mutations with different ∆∆G values. We have used it here to compare against the performance of previous methods, which were applied to this dataset [11, 14, 15].

These limitations prompted us to propose a new experimental antibody-antigen ∆∆G bench-marking dataset, Experimental_∆∆G_608 (Supplementary Table 1), consisting of 608 single-point mutations, obtained by rigorous filtering of the SKEMPI 2.0 database [24]. We filtered out non-binder mutations, mutations with imprecisely measured affinity and duplicated mutations. A more detailed description of the datasets and further information on the filtering protocol is included in the Supplementary Information.

### Synthetic ∆∆G data preparation

To investigate affinity prediction without the constraint of dataset size, we generated a synthetic dataset orders of magnitude larger than the experimental datasets (Figure 1b).

We downloaded structurally resolved antibody-protein antigen complexes from SAbDab [21, 22], resulting in 6077 non-redundant entries from 3065 PDB files (SAbDab accession date: 1 9 May 2022). Twenty-seven PDBs with only C_*α*_ residues resolved were removed from the dataset. We renumbered the PDB files using a custom script, to prevent issues with insertion numbering in subsequent steps with FoldX, and repaired the PDBs using FoldX RepairPDB [8]. We removed from the dataset 25 PDBs for which the repair did not run to completion. We then clustered the antibody-antigen complexes based on a 90% length-matched CDR sequence identity threshold (see below), resulting in 1475 clusters. One complex per cluster was carried forward for exhaustive interface mutagenesis: all interface residues, defined as being within 4 Å of the binding partner, were mutated to every other amino acid using FoldX BuildModel [8]. The Interaction Energy for each WT and mutant complex was estimated with FoldX AnalyseComplex [8], and the FoldX ∆∆G determined as:

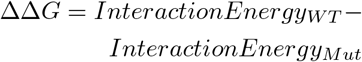

such that a negative ∆∆G represents a destabilizing mutation.

We excluded mutations where the WT amino acid was ‘X’, the chain identifier was a number, the antibody and antigen were *>*4 Å apart and the FoldX Interaction Energy calculation failed. The final dataset (Synthetic_∆∆G_942723, Supplementary Table 1) consisted of 942,723 mutations from 1471 antibody-antigen complexes from 1409 PDBs. The dataset and download links can be found at https://github.com/oxpig/Graphinity.

### Train-validation-test cutoffs

We numbered all antibody sequences with ANARCI [29] using the IMGT numbering scheme [30]. The CDRs were extracted, concatenated and binned based on length. We applied CD-HIT [31], with varying sequence identity cutoffs, to cluster the length-matched CDRs. Seventy percent was the lowest threshold rounded to 10 for which CD-HIT ran (for CDRs and antigen sequences).

The AB-Bind Experimental_∆∆G_645 dataset contained non-antibody-antigen complexes, which could not be clustered by CDR sequence identity. The sequences of each of the chains in these complexes had less than 90% sequence identity with each other and each complex was considered as its own cluster.

We generated a synthetic dataset imposing an antigen sequence identity cutoff in addition to the antibody CDR sequence identity cutoff. In this case, antigen sequences were extracted from the PDB structures using the Bio.PDB.PDBParser module and clustered using CD-HIT with a 70% sequence identity cutoff. Clusters from the antibody CDR- and antigen-based sequence identity cutoffs were merged such that no cluster had a complex with *>*70% length-matched CDR sequence identity to an antibody in another cluster nor *>*70% sequence identity to an antigen in another cluster.

Train-validation-test datasets were generated with an 80%-10%-10% split, with respect to the full dataset size. The datasets were sampled such that no cluster had members in more than one dataset, with the exception of datasets split with no cutoff. For 10-fold cross-validation, we generated ten dataset folds using the CD-HIT clusters, such that no cluster had members in more than one fold.

Unless otherwise specified, models were built from a single-fold 80%-10%-10% split with a 90% length-matched CDR sequence identity cutoff.

### Varying synthetic dataset amounts

To investigate the role of dataset size on model performance, we trained models on a subset of the full, large-scale synthetic dataset (Synthetic_∆∆G_942723). These subsets were randomly sampled from the respective train and validation datasets (Synthetic_∆∆G_{580-450000}, Supplementary Table 1). All models were evaluated on the same test set, consisting of 94,126 mutations (one fold, held-out test set). A 90% length-matched CDR sequence identity cutoff was applied between respective train, validation and test sets.

### Varying synthetic dataset diversity

We explored the importance of dataset diversity, in addition to dataset size, for model performance via the following three metrics:

- The number of antibody clusters, following clustering with a 90% length-matched CDR sequence identity cutoff
- The number of amino acid substitution types (e.g. Arg to Lys; Arg to Ala)
- The distribution of amino acid substitutions in the complex: mutation locations were classified based on binding partner (antibody/antigen) and proximity to the interface center; for the latter, the interface was divided into two areas (inner and outer shell) defined by concentric circles where, assuming that the interface is approximately flat, the outer shell circle was defined with a radius 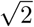 times the radius of the inner shell circle, to produce two equal areas)

We generated training and validation datasets minimizing and maximizing the different metrics of diversity (Synthetic_∆∆G_100000_{sequence/substitution_type/substitution_distribution}_{min/max}, Supplementary Table 1). The test data was kept the same in each case. The respective training, validation and test datasets consisted of 100,000 mutations combined. A 90% length-matched CDR sequence identity cutoff was applied between each.

### Investigating model robustness to noise

We assessed the robustness of our models to noise by (1) shuffling and (2) applying random noise from a Gaussian distribution to the training and validation dataset affinity labels. In each of these cases, the test data remained unmodified.

- **Shuffling:** Varying percentages of the training and validation ∆∆G dataset labels were shuffled. The effective shuffling percentage was not necessarily equal to the percentage of the dataset which was shuffled, as some labels are the same and others were shuffled back into the same place.
- **Gaussian noise:** Gaussian noise was applied by adding random values generated from a normal distribution, using numpy.normal, with a set scale (0.5, 1, 2, 5 or 10) to the training and validation datasets.

### Graphinity: Equivariant Graph Neural Network architecture

We developed a deep learning EGNN architecture to predict change in antibody-antigen binding affinity (Figure 1a). Our model is composed of three E(n) Equivariant Graph Convolutional (EGC) layers [20] with a hidden dimension of 128. The model takes the 3D coordinates of a protein complex structure (PDB file) as input and generates an atomic-resolution graph with nodes representing non-hydrogen atoms and edges representing interactions between nodes *<*4 Å apart. The node features are a one-hot encoded vector describing the LibMolGrid atom type [32] and the edge features a one-hot encoded vector describing whether the edge is intra-binding partner, i.e. between atoms on the same binding partner, or inter-binding partner, i.e. between atoms on different binding partners. The graphs represent the mutation site neighborhood (for ∆∆G prediction: atoms on the same chain as the mutated residue within 4 Å of the mutated residue (local neighborhood) and atoms on the binding partner chain within 4 Å of these local neighborhood atoms). The models were trained with Mean Squared Error loss. The architecture was implemented using PyTorch and PyTorch Geometric.

Models were trained using PyTorch Lightning for 500 epochs, with the exception of the synthetic ∆∆G dataset models, which, due to the high computational costs, were trained for 10 epochs. Model parameters and training times are included in the Supplementary Information.

For ∆∆G prediction, we generated and aggregated graphs of the WT and mutant structures. Both graphs were fed through the three E(n) EGC layers and the resulting embeddings subtracted from one another (*WT −Mutant*) prior to the last linear layer.

In the models generated with datasets limited to a specific amino acid substitution and transfer learning, we initialized model weights with those from the model trained on the full dataset. In these cases, the learning rate was set to 0.0001.

### Tree-based model trained on featurized structures

To investigate the role of model architecture, we generated a tree-based model trained on featurized structures. Features were derived from the antibody-antigen structures as in mCSM-AB2 [15]: FoldX AnalyseComplex energetic terms [8], Arpeggio interactions [33], pharmacophore vectors [34], buried surface area [35] and Position Specific Scoring Matrix evolutionary term [36] (more detail in the Supplementary Information). We generated an Extra Trees model with 300 estimators as in mCSM-AB2. This is not a direct comparison to the mCSM-AB2 model, as we do not incorporate the graph-based features of the CSM-based models.

We applied this featurization and subsequent Extra Trees model to the Experimental_∆∆G_608 dataset. Given the time required for featurization, it was computationally infeasible to apply this approach to the large synthetic dataset.

### Trastuzumab variants

We obtained the dataset of Trastuzumab CDRH3 variants and corresponding binary binding labels from [12]. The sequences were mutated at 10 amino acid positions in the CDRH3 [12]. The variants which had been labeled as both binding and non-binding were assigned the binding label, as in [12]. This resulted in 36,391 variants, 11,277 of which were labeled as binding. We split the dataset (1) randomly using sklearn.model_selection.train_test_split and (2) with a clonotype plus sequence-identity split. For (2), variants were clustered based on the V- and J-gene assignments, as labeled by ANARCI [29], and sequence identity of the CDRH3 (limited to the 10 mutated positions). Sequence identity in this case describes the maximum allowed edit distance from a representative sequence (cluster center). For example, a minimum identity of 70% allows edit distances of up to three residues from the cluster center. We used the clonotype and sequence identity approach as CD-HIT did not run with the 10-position variant sequences due to their short length.

The Trastuzumab datasets were prepared with a 70%-15%-15% train-validation-test split to allow comparison with [12].

We modeled structures of the Trastuzumab variants in complex with HER2 using the FoldX BuildModel function starting from a FoldX-‘repaired’ structure of PDB 1N8Z [8, 37].

We adapted the Graphinity architecture for this task. The input was changed to be one graph only (and subsequently there was no subtraction of embeddings before the final layer) and the graphs were formed from the 10 mutated CDRH3 residues and surrounding neighbourhood (antibody atoms within 4 Å of CDRH3 atoms (antibody neighborhood), antigen atoms within 4 Å of the antibody neighborhood and antigen atoms within 4 Å of these antigen atoms). We also updated the model for classification by changing the loss function (to Binary Cross-Entropy with Logits) and accuracy metrics (to ROC AUC and AP).

### Correlations

The Pearson’s and Spearmans’ rank correlations were calculated using scipy.stats.pearsonr and scipy.stats.spearmanr, respectively.

## Data visualization

Figures were generated using matplotlib, seaborn and PowerPoint. Colors were selected in part using ColorBrewer 2.0 (https://colorbrewer2.org).

## Data availability

The synthetic ∆∆G datasets and links to download the corresponding PDBs can be found at https://github.com/oxpig/Graphinity.

## Code availability

The Graphinity EGNN model code is available at https://github.com/oxpig/Graphinity.

## Author contributions

A.M.H. and C.S. developed the code. A.M.H. and C.M.D. designed the experiments. A.M.H. performed the experiments and analyses. L.C. prepared the Trastuzumab data. A.M.H. wrote the manuscript and all other authors revised it. C.M.D. supervised the work. All authors read and approved the final version of the manuscript.

## References

[1] Makowski, E. K. et al. Reduction of therapeutic antibody self-association using yeast-display selections and machine learning. mAbs 14, 2146629 (2022).

[2] Marks, C., Hummer, A. M., Chin, M. & Deane, C. M. Humanization of antibodies using a machine learning approach on large-scale repertoire data. Bioinformatics 37, 4041–4047 (2021).

[3] Prihoda, D. et al. Biophi: A platform for antibody design, humanization, and humanness evaluation based on natural antibody repertoires and deep learning. mAbs 14, 2020203 (2022).

[4] Makowski, E. K. et al. Co-optimization of therapeutic antibody affinity and specificity using machine learning models that generalize to novel mutational space. Nature Communications 13 (2022).

[5] Harvey, E. P. et al. An in silico method to assess antibody fragment polyreactivity. Nature Communications 13, 7554 (2022).

[6] Wang, B., Gallolu Kankanamalage, S., Dong, J. & Liu, Y. Optimization of therapeutic antibodies. Antibody Therapeutics 4, 45–54 (2021).

[7] Jarmoskaite, I., AlSadhan, I., Vaidyanathan, P. P. & Herschlag, D. How to measure and evaluate binding affinities. eLife 9, e57264 (2020).

[8] Schymkowitz, J. et al. The FoldX web server: An online force field. Nucleic Acids Research 33, 382–388 (2005).

[9] Barlow, K. A. et al. Flex ddG: Rosetta Ensemble-Based Estimation of Changes in Protein-Protein Binding Affinity upon Mutation. Journal of Physical Chemistry B 122, 5389–5399 (2018).

[10] Leman, J. K. et al. Macromolecular modeling and design in Rosetta: recent methods and frameworks. Nature Methods 17, 665–680 (2020).

[11] Pires, D. E. & Ascher, D. B. mCSM-AB: a web server for predicting antibody–antigen affinity changes upon mutation with graph-based signatures. Nucleic Acids Research 44, W469–W473 (2016).

[12] Mason, D. M. et al. Optimization of therapeutic antibodies by predicting antigen specificity from antibody sequence via deep learning. Nature Biomedical Engineering 5, 600–612 (2021).

[13] Bachas, S. et al. Antibody optimization enabled by artificial intelligence predictions of binding affinity and naturalness. bioRxiv 2022.08.16.504181 (2022).

[14] Wang, M., Cang, Z. & Wei, G.-w. A topology-based network tree for the prediction of protein–protein binding affinity changes following mutation. Nature Machine Intelligence 2, 116–123 (2020).

[15] Myung, Y., Rodrigues, C. H., Ascher, D. B. & Pires, D. E. mCSM-AB2: Guiding rational antibody design using graph-based signatures. Bioinformatics 36, 1453–1459 (2020).

[16] Geng, C., Xue, L. C., Roel-Touris, J. & Bonvin, A. M. J. J. Finding the g spot: Are predictors of binding affinity changes upon mutations in protein–protein interactions ready for it? WIREs Computational Molecular Science 9, e1410 (2019).

[17] Sirin, S., Apgar, J. R., Bennett, E. M. & Keating, A. E. AB-Bind: Antibody binding mutational database for computational affinity predictions. Protein Science 25, 393–409 (2016).

[18] Liu, X., Luo, Y., Li, P., Song, S. & Peng, J. Deep geometric representations for modeling effects of mutations on protein-protein binding affinity. PLOS Computational Biology 17, 1–28 (2021).

[19] Behbahani, Y. M., Laine, E. & Carbone, A. Deep local analysis estimates effects of mutations on protein-protein interactions. bioRxiv (2022).

[20] Satorras, V. G., Hoogeboom, E. & Welling, M. E(n) equivariant graph neural networks. arXiv (2021).

[21] Dunbar, J. et al. SAbDab: The structural antibody database. Nucleic Acids Research 42, 1140–1146 (2014).

[22] Schneider, C., Raybould, M. I. J. & Deane, C. M. SAbDab in the age of biotherapeutics: updates including SAbDab-nano, the nanobody structure tracker. Nucleic Acids Research 50, D1368–D1372 (2021).

[23] Li, Y., Huang, Y., Swaminathan, C. P., Smith-Gill, S. J. & Mariuzza, R. A. Magnitude of the hydrophobic effect at central versus peripheral sites in protein-protein interfaces. Structure 13, 297–307 (2005).

[24] Jankauskaite, J., Jiménez-García, B., Dapkunas, J., Fernández-Recio, J. & Moal, I. H. SKEMPI 2.0: An updated benchmark of changes in protein-protein binding energy, kinetics and thermodynamics upon mutation. Bioinformatics 35, 462–469 (2019).

[25] Olsen, T. H., Abanades, B., Moal, I. H. & Deane, C. M. Ka-search: Rapid and exhaustive sequence identity search of known antibodies. bioRxiv 2022.11.01.513855 (2022).

[26] Rosace, A. et al. Automated optimisation of solubility and conformational stability of antibodies and proteins. bioRxiv (2022).

[27] Van Durme, J. et al. A graphical interface for the FoldX forcefield. Bioinformatics 27, 1711–1712 (2011).

[28] Bernett, J., Blumenthal, D. B. & List, M. Cracking the black box of deep sequence-based protein-protein interaction prediction. bioRxiv (2023).

[29] Dunbar, J. & Deane, C. M. ANARCI: Antigen receptor numbering and receptor classification. Bioinformatics 32, 298–300 (2016).

[30] Lefranc, M.-P. et al. Imgt unique numbering for immunoglobulin and t cell receptor variable domains and ig superfamily v-like domains. Developmental Comparative Immunology 27, 55–77 (2003).

[31] Fu, L., Niu, B., Zhu, Z., Wu, S. & Li, W. CD-HIT: Accelerated for clustering the next-generation sequencing data. Bioinformatics 28, 3150–3152 (2012).

[32] Sunseri, J. & Koes, D. R. libmolgrid: Graphics processing unit accelerated molecular gridding for deep learning applications. Journal of Chemical Information and Modeling 60, 1079–1084 (2020).

[33] Jubb, H. C. et al. Arpeggio: A web server for calculating and visualising interatomic interactions in protein structures. Journal of Molecular Biology 429, 365–371 (2017).

[34] Pires, D. E., Ascher, D. B. & Blundell, T. L. mCSM: Predicting the effects of mutations in proteins using graph-based signatures. Bioinformatics 30, 335–342 (2014).

[35] Lee, B. & Richards, F. M. The interpretation of protein structures: Estimation of static accessibility. Journal of Molecular Biology 55 (1971).

[36] Altschul, S. F. et al. Gapped BLAST and PSI-BLAST: a new generation of protein database search programs. Nucleic Acids Research 25, 3389–3402 (1997).

[37] Cho, H.-S. et al. Structure of the extracellular region of her2 alone and in complex with the herceptin fab. Nature 421, 756–760 (2003).

