## Supplementary Information for "Investigating the Volume and Diversity of Data Needed for Generalizable Antibody-Antigen ∆∆G Prediction"

#### Supplementary Tables and Figures

Supplementary Table 1: Descriptions of the experimental and synthetic $\Delta\Delta G$ datasets to which Graphinity was applied. Ab: antibody, Ag: antigen, AA: amino acid. For definitions of inner and outer shell see Methods.

| Dataset Name | Experimental/ Synthetic | Description | | Train-Val-Test Split (CDR Sequence Identity Cutoff) | Number of Mutations | Number of Complexes | Number of AA Substitution Types | Mutation Distribution (# of muts. in Ab Inner, Ab Outer, Ag Inner, Ag Outer) |
|---|---|---|---|---|---|---|---|---|
| Experimental_ΔΔG_645 – Reverse Mutations + Non-Binders | Experimental | Dataset of single-point mutations from AB-Bind (Sirin et al., 2016) | – reverse mutations, + non-binders | None (Random), 100%, 90%, 70% | 645 | 24 (plus 5 homology models) | 141 | 185, 148, 172, 140 |
| Experimental_ΔΔG_645 – Reverse Mutations – Non-Binders | | | – reverse mutations, – non-binders | | 618 | | 136 | 170, 136, 172, 140 |
| Experimental_ΔΔG_645 + Reverse Mutations + Non-Binders | | | + reverse mutations, +non-binders | | 1,290 | | 224 | 370, 296, 344, 280 |
| Experimental_ΔΔG_645 + Reverse Mutations – Non-Binders | | | + reverse mutations, – non-binders | | 1,236 | | 216 | 340, 272, 344, 280 |
| Experimental_ΔΔG_608 | | Dataset of single-point mutations filtered from SKEMPI 2.0 (Jankauskaite et al., 2019) | – reverse mutations | | 608 | 33 | 163 | 232, 138, 151, 87 |
| Experimental_ΔΔG_608 + Reverse Mutations | | | + reverse mutations | | 1,216 | | 232 | 464, 276, 302, 174 |
| Synthetic_ΔΔG_ 942723 | Synthetic | Synthetic single-point mutation ΔΔG data generated using FoldX | | None (Random), 100%, 90%, 70%, 70% + 70% Ag seq. identity cutoff | 942,723 | 1,471 | 380 | 326,990, 155,439, 328,130, 132,164 |
| Synthetic_ΔΔG_ 942723_shuffled | | Synthetic_ΔΔG_942723 with a percentage of ΔΔG labels shuffled (i.e. incorrect) | | 90% | | | | |
| Synthetic_ΔΔG_ 942723_gaussian_ noise | | Synthetic_ΔΔG_942723 with random noise sampled from Gaussian distributions with varying scales added | | | | | | |
| Synthetic_ΔΔG_580 | | Train + validation datasets of varying sizes randomly sampled from the respective Synthetic_ΔΔG_942723 datasets | | | 580 | 462 | 269 | 200, 87, 212, 81 |
| Synthetic_ΔΔG_900 | | | | | 900 | 645 | 313 | 296, 137, 331, 136 |
| Synthetic_ΔΔG_4500 | | | | | 4,500 | 1,264 | 377 | 1,519, 711, 1,602, 668 |
| Synthetic_ΔΔG_9000 | | | | | 9,000 | 1,316 | 380 | 3,052, 1,468, 3,180, 1,300 |
| Synthetic_ΔΔG_45000 | | | | | 45,000 | 1,324 | 380 | 15,522, 7,443, 15,843, 6,192 |
| Synthetic_ΔΔG_90000 | | | | | 90,000 | 1,324 | 380 | 31,337, 14,800, 31,387, 12,476 |
| Synthetic_ΔΔG_450000 | | | | | 450,000 | 1,324 | 380 | 156,449, 74,016, 156,945, 62,590 |
| Synthetic_ΔΔG_848597 | | | | | 848,597 | 1,324 | 380 | 294,614, 139,802, 296,305, 117,876 |
| Synthetic_ΔΔG_94126 | | Test dataset to which models trained on the train + validation datasets of varying sizes were applied | | | 94,126 | 147 | 380 | 32,376, 15,637, 31,825, 14,288 |
| Synthetic_ΔΔG_ 100000_sequence_min | | Datasets of 90,000 mutations sampled from Synthetic_ΔΔG_ 942723 to: [NB train and validation datasets only, which are 80,000 and 10,000 mutations respectively] | minimize antibody CDR sequence diversity | | Train: 80,000; Val: 10,000 | 86 (Train + Val) | 380 | 34,143, 10,678, 32,785, 12,394 |
| Synthetic_ΔΔG_ 100000_sequence_max | | | maximize antibody CDR sequence diversity | | | 1,324 (Train + Val) | 380 | 30,744, 15,420, 31,172, 12,664 |
| Synthetic_ΔΔG_ 100000_substitution_ type_min | | | minimize antibody substitution type diversity | | | 1,293 (Train + Val) | 16 | 48,747, 20,314, 14,940, 5,999 |
| Synthetic_ΔΔG_ 100000_substitution_ type_max | | | maximize antibody substitution type diversity | | | 1,324 (Train + Val) | 380 | 35,549, 12,464, 27,987, 14,000 |
| Synthetic_ΔΔG_ 100000_mutation_ distribution_min | | | minimize antibody mutation distribution diversity | | | 1,321 (Train + Val) | 380 | 0, 0, 90,000, 0 |
| Synthetic_ΔΔG_ 100000_mutation_ distribution_max | | | maximize antibody mutation distribution diversity | | | 1,324 (Train + Val) | 380 | 22,505, 22,498, 22,499, 22,498 |
| Synthetic_ΔΔG_ 100000_randomly_ sampled | | Train and validation datasets randomly sampled from Synthetic_ΔΔG_ 942723, for which no complex overlaps with any in Synthetic_ΔΔG_100000_diversity_test_set | | | | 1,332 (Train + Val) | 380 | 31,283, 14,800, 31,345, 12,572 |
| Synthetic_ΔΔG_ 100000_diversity_ test_set | | Test dataset for all train/val diversity datasets – consists of 10,000 mutations, for which no complex overlaps with any in the train and validation sets | | | Test: 10,000 | 139 (Test) | 380 | 3,468, 3,377, 1,763, 1,392 |

Supplementary Table 2: Pharmacophore counts for each amino acid, as used in the tree-based model featurization.

| AA | Neutral | H-bond Donor | H-bond Acceptor | Hydro-phobic | Aromatic | Positive | Negative | Sulphur |
|---|---|---|---|---|---|---|---|---|
| **A** | 2 | 1 | 1 | 1 | 0 | 0 | 0 | 0 |
| **C** | 2 | 1 | 1 | 1 | 0 | 0 | 0 | 1 |
| **D** | 3 | 1 | 3 | 1 | 0 | 0 | 2 | 0 |
| **E** | 3 | 1 | 3 | 2 | 0 | 0 | 2 | 0 |
| **F** | 2 | 1 | 1 | 7 | 6 | 0 | 0 | 0 |
| **G** | 2 | 1 | 1 | 0 | 0 | 0 | 0 | 0 |
| **H** | 2 | 3 | 3 | 6 | 5 | 2 | 0 | 0 |
| **I** | 2 | 1 | 1 | 4 | 0 | 0 | 0 | 0 |
| **K** | 2 | 2 | 1 | 4 | 0 | 1 | 0 | 0 |
| **L** | 2 | 1 | 1 | 4 | 0 | 0 | 0 | 0 |
| **M** | 2 | 1 | 1 | 4 | 0 | 0 | 0 | 0 |
| **N** | 3 | 2 | 2 | 1 | 0 | 0 | 0 | 0 |
| **P** | 1 | 0 | 1 | 5 | 0 | 0 | 0 | 0 |
| **Q** | 3 | 2 | 2 | 2 | 0 | 0 | 0 | 0 |
| **R** | 3 | 4 | 1 | 3 | 0 | 3 | 0 | 0 |
| **S** | 2 | 2 | 2 | 1 | 0 | 0 | 0 | 0 |
| **T** | 2 | 2 | 2 | 2 | 0 | 0 | 0 | 0 |
| **V** | 2 | 1 | 1 | 3 | 0 | 0 | 0 | 0 |
| **W** | 2 | 2 | 1 | 10 | 9 | 0 | 0 | 0 |
| **Y** | 2 | 2 | 2 | 7 | 6 | 0 | 0 | 0 |

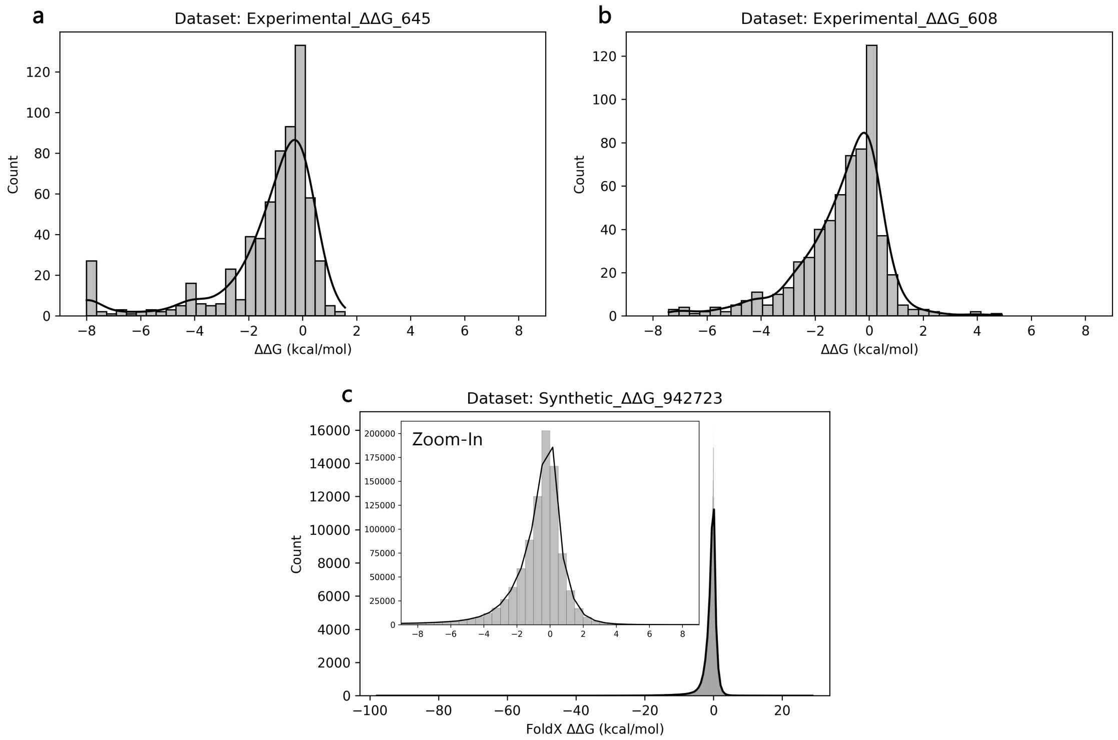


Supplementary Figure 1: The distributions of the $\Delta\Delta G$ values of the base datasets used in this study: (a) Experimental_ΔΔG_645, (b) Experimental_ΔΔG_608, (c) Synthetic_ΔΔG_942723. The solid lines are kernel density estimates.

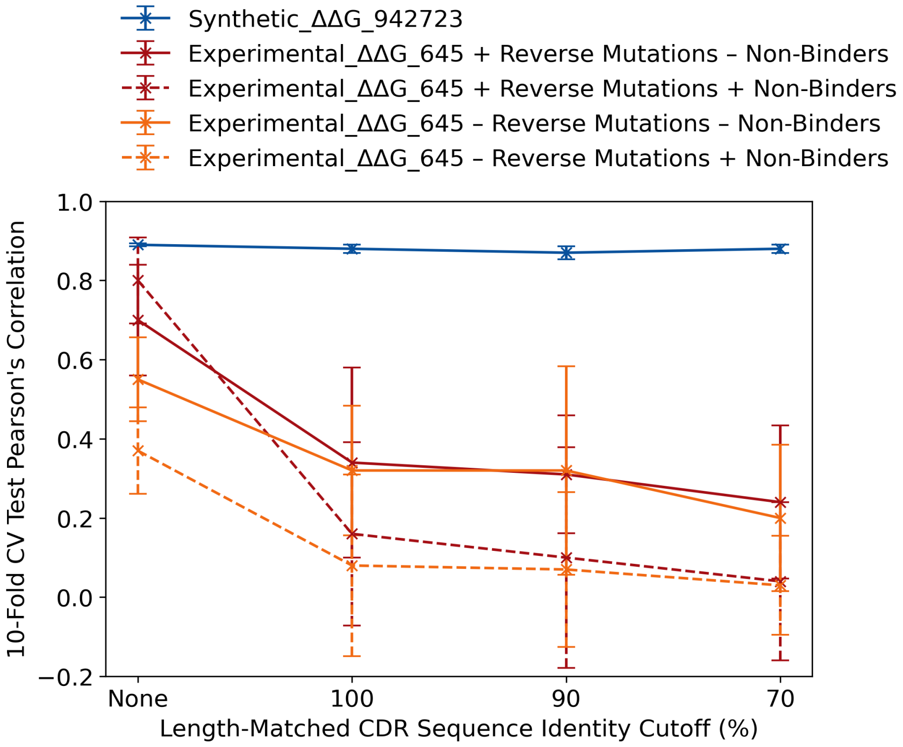


Supplementary Figure 2: The Pearson's correlations of Graphinity on different train-validation-test cutoffs applied to the Experimental_ΔΔG_645 dataset (red, orange) and Synthetic_ΔΔG_942723 dataset (blue). This is Figure 2b including error bars, which represent the standard deviation in Pearson's correlation across the 10 folds of 10-fold cross-validation (CV).

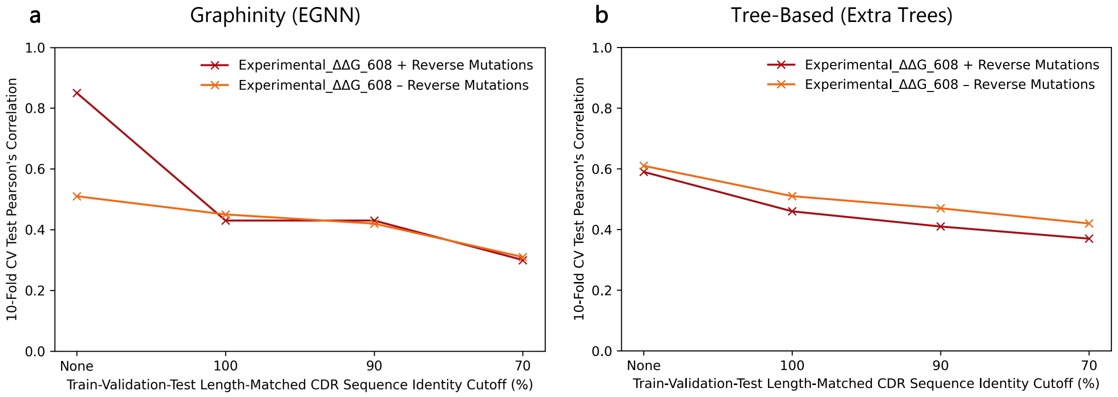


Supplementary Figure 3: Performance of (a) Graphinity (EGNN architecture) and (b) a tree-based (Extra Trees) model on the Experimental_ΔΔG_608 dataset, with and without reverse mutations, at different length-matched CDR sequence identity cutoffs.

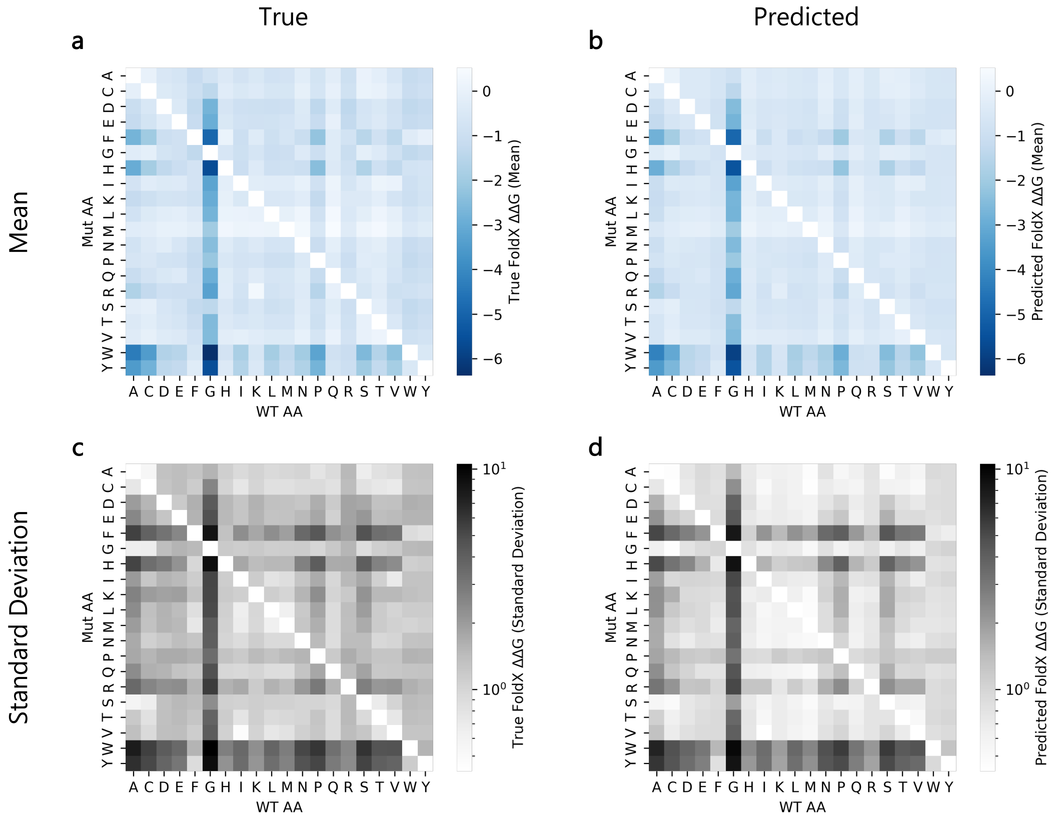


Supplementary Figure 4: (a,c) ΔΔG values from the Synthetic_ΔΔG_942723 dataset, separated by amino acid substitution. (b,d) ΔΔG values predicted by Graphinity applied to the Synthetic_ΔΔG_942723 dataset (10-fold cross-validation, 90% length-matched CDR sequence identity cutoff), separated by amino acid substitution. Top row: mean, bottom row: standard deviation. WT: wild-type, Mut: mutant.

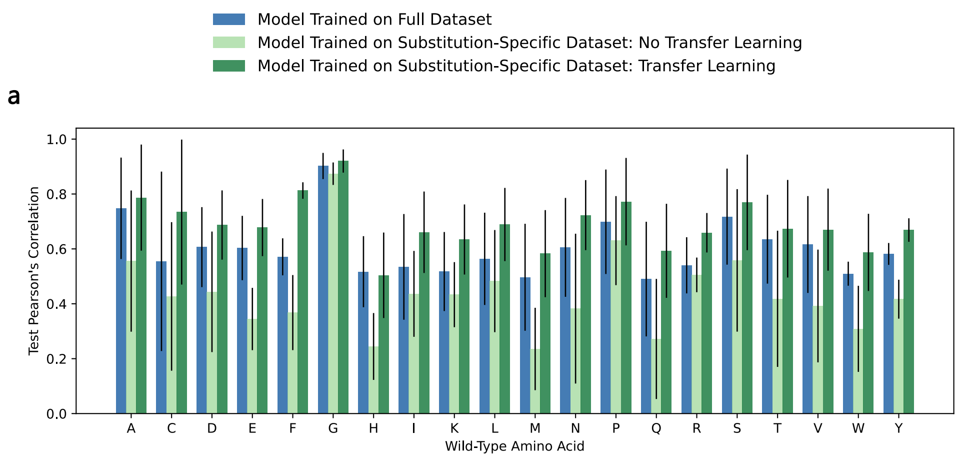


Supplementary Figure 5: Graphinity performance on amino acid substitutions when trained on the full dataset (blue), substitution-specific dataset (light green) and substitution-specific dataset with weights initialized from the model trained on the full dataset (dark green). The results were grouped and averaged by (a) wild-type amino acid and (b) mutant amino acid. The error bars represent the standard deviation.

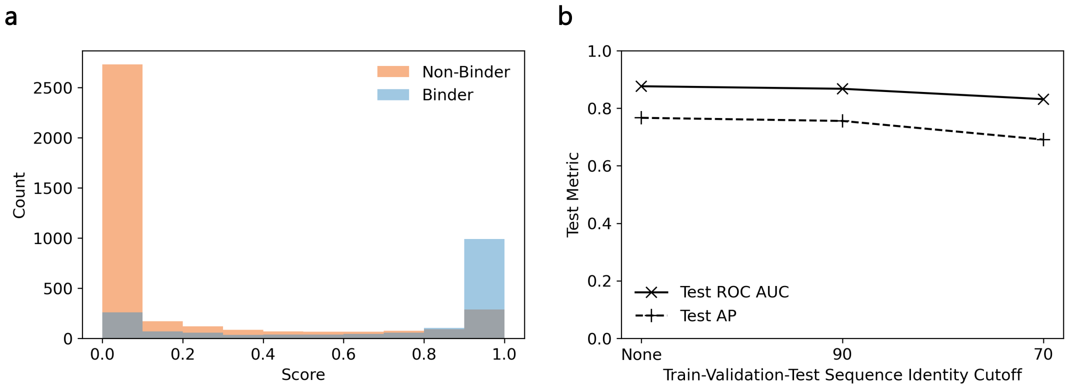


Supplementary Figure 6: (a) Graphinity scoring of 36,391 Trastuzumab CDRH3 variants [1] (randomly split data). (b) Model performance with clonotype and sequence identity cutoffs imposed between the train, validation and test datasets. In cases where a sequence identity cutoff (value shown on the x-axis) was applied, the data was also separated by clonotype (V- and J-gene assignments).

### Supplementary Methods

#### Filtering SKEMPI v2.0

We downloaded the SKEMPI 2.0 database, which contains data on changes to the binding affinity of structurally resolved protein-protein interactions in response to mutations [2]. The total database consists of 7085 entries, 1150 of which are for antibody-antigen interactions. We removed non-antibody-antigen complexes and further filtered the antibody-antigen dataset by removing multi-point mutations, non-binder mutations, mutations in which the affinity could not be measured exactly and duplicates of mutations. When removing duplicate mutations, we preferentially retained those with kinetic data, based on the measurement method (SPR > ITC > KinExA > FL > IASP > SP > CSPRIA > ELISA > BI) and based on the temperature (298 > 296 > 303 > 310 > 283 > 298(assumed)). The final filtered dataset contained 608 single-point mutations from 44 complexes.

We calculated ΔΔG from the SKEMPI 2.0 database as:

$$\Delta G = RT * ln(K_d)$$

$$\Delta\Delta G = \Delta G_{WT} - \Delta G_{Mutant}$$

#### Tree-based ΔΔG prediction

##### Feature extraction

The approach for featurization was inspired by the mCSM-AB2 method [3]. For each feature, we calculated the difference between the values for the WT and mutant structures (WT − Mutant).

- FoldX AnalyseComplex energetic terms: We used the FoldX AnalyseComplex function [4] to calculate interaction energetic terms for the WT and mutant complexes.
- Arpeggio interactions: We calculated the inter-protein interface interactions (e.g. H-bonds and ionic interactions) of the complexes using Arpeggio [5].
- Pharmacophore vectors: To represent the change in amino acid upon mutation, we calculated a change in the pharmacophore counts, adapted from [6]. We assigned pharmacophores (e.g. hydrophobic, H-bond acceptor or H-bond donor) to each atom in each amino acid and took a sum across the amino acid (Supplementary Table 2). To note, an atom can have more than one pharmacophore.
- Buried surface area: We calculated the buried surface area (BSA) for each binding partner (antibody and antigen) in each complex using the PSA program [7]: $BSA = SA_{free} - SA_{bound}$ An average change in BSA across the two binding partners was calculated.
- Position Specific Scoring Matrix evolutionary term: A measure of residue conservation at a position was captured in Position Specific Scoring Matrices (PSSMs). We calculated the PSSM scores using PSI-BLAST [8] (parameters: evolutionary scoring matrix = PAM30, num_iterations = 3, evalue = 1E-10, seg = Yes, comp_based_stats = 1, and db = swissprot) as in [3].

##### Model development

We generated ExtraTrees regression models using the Python scikit-learn ExtraTreesRegressor package with 300 estimators (as in [3]) and remaining default parameters.

#### Graphinity architecture

##### Parameters

The model parameters were set as:

Optimizer: Adam
Learning rate: 0.001
Batch size: 32
Dropout: 0.2
Weight decay: 1e-16
Graph readout: global_max_pool over nodes
Tanh activation at the output of the coordinate function: True
Update coords: True

##### Model training times

The following model training times are given for training with 1 GPU (NVIDIA RTX 6000) and 4 CPUs on a single data fold (80/10/10 train-validation-test data split).

Experimental_ΔΔG_645 (500 epochs): ca. 1 hour

Experimental_ΔΔG_608 (500 epochs): ca. 1 hour

Synthetic_ΔΔG_942723 (10 epochs): ca. 19.5 hours

Trastuzumab Variants (500 epochs): ca. 35 hours

#### Evolutionarily grounded mutations

A recent study demonstrated that the likelihood of FoldX incorrectly predicting a mutation to be stabilizing (in this case, independent of an antigen) could be decreased by up to 11% by limiting FoldX predictions to mutations that are observed naturally [9]. We investigated the effect of limiting our test dataset to such 'evolutionarily grounded' mutations, as defined in [9], on model performance.

##### Identifying evolutionarily grounded mutations

We obtained the Position Specific Scoring Matrices (PSSMs) generated from subsets of the Observed Antibody Space database [10, 11, 12] and corresponding custom code for calculating log-likelihoods from the authors of [9]. As in [9], we defined the 'evolutionarily grounded' mutations as those with a positive log-likelihood and which have a log-likelihood greater than is seen for the wild-type residue [9].

We mapped the log-likelihood scores to the dataset mutations via the Aho numbering scheme [13] used for the PSSMs, with sequences numbered using ANARCI [14]. There were 10 PDBs where ANARCI failed to number a chain with the Aho numbering scheme (3U2S, 4DQO, 4Y5Y, 6BPE, 6E1K, 6OPA, 6U0N, 7EY0, 7LF8, 7LY9). We applied this approach to mutations from antibody chains from humans or mice, as identified in SAbDab [15, 16], as the PSSMs were restricted to these species.

Complexes with human or mouse antibodies made up 75% percent (710,562 mutations) of the full synthetic dataset. Just over half of these (52%, 366,862) were for mutations to an antibody chain. The final 'evolutionarily grounded' dataset consisted of 47,983 mutations.

##### Model performance on evolutionarily grounded mutations

The performance of our model was stable on a test dataset limited to 'evolutionarily grounded' mutations from human and mouse sequences (see Methods), with a Pearson's correlation of 0.89. Conversely, Graphinity also performed well (Pearson's correlation = 0.85) on non-evolutionarily grounded mutations from human and mouse sequences.

### Supplementary Information References


[1] Mason, D. M. *et al.* Optimization of therapeutic antibodies by predicting antigen specificity from antibody sequence via deep learning. *Nature Biomedical Engineering* **5**, 600–612 (2021).

[2] Jankauskaite, J., Jiménez-García, B., Dapkunas, J., Fernández-Recio, J. & Moal, I. H. SKEMPI 2.0: An updated benchmark of changes in protein-protein binding energy, kinetics and thermodynamics upon mutation. *Bioinformatics* **35**, 462–469 (2019).

[3] Myung, Y., Rodrigues, C. H., Ascher, D. B. & Pires, D. E. mCSM-AB2: Guiding rational antibody design using graph-based signatures. *Bioinformatics* **36**, 1453–1459 (2020).

[4] Schymkowitz, J. *et al.* The FoldX web server: An online force field. *Nucleic Acids Research* **33**, 382–388 (2005).

[5] Jubb, H. C. *et al.* Arpeggio: A Web Server for Calculating and Visualising Interatomic Interactions in Protein Structures. *Journal of Molecular Biology* **429**, 365–371 (2017).

[6] Pires, D. E., Ascher, D. B. & Blundell, T. L. mCSM: Predicting the effects of mutations in proteins using graph-based signatures. *Bioinformatics* **30**, 335–342 (2014).

[7] Lee, B. & Richards, F. M. The interpretation of protein structures: Estimation of static accessibility. *Journal of Molecular Biology* **55** (1971).

[8] Altschul, S. F. *et al.* Gapped BLAST and PSI-BLAST: a new generation of protein database search programs. *Nucleic Acids Research* **25**, 3389–3402 (1997).

[9] Rosace, A. *et al.* Automated optimisation of solubility and conformational stability of antibodies and proteins. *bioRxiv* (2022).

[10] Prihoda, D. *et al.* BioPhi: A platform for antibody design, humanization, and humanness evaluation based on natural antibody repertoires and deep learning. *mAbs* **14** (2022).

[11] Olsen, T. H., Boyles, F. & Deane, C. M. Observed antibody space: A diverse database of cleaned, annotated, and translated unpaired and paired antibody sequences. *Protein Science* **31**, 141–146 (2022).

[12] Kovaltsuk, A. *et al.* Observed Antibody Space: A Resource for Data Mining Next-Generation Sequencing of Antibody Repertoires. *The Journal of Immunology* **201**, 2502–2509 (2018).

[13] Honegger, A. & Plückthun, A. Yet another numbering scheme for immunoglobulin variable domains: An automatic modeling and analysis tool. *Journal of Molecular Biology* **309**, 657–670 (2001).

[14] Dunbar, J. & Deane, C. M. ANARCI: Antigen receptor numbering and receptor classification. *Bioinformatics* **32**, 298–300 (2016).

[15] Dunbar, J. *et al.* SAbDab: The structural antibody database. *Nucleic Acids Research* **42**, 1140–1146 (2014).

[16] Schneider, C., Raybould, M. I. J. & Deane, C. M. SAbDab in the age of biotherapeutics: updates including SAbDab-nano, the nanobody structure tracker. *Nucleic Acids Research* **50**, D1368–D1372 (2021).
